# Selectivity and ranking of tight-binding JAK-STAT inhibitors using Markovian milestoning with Voronoi tessellations

**DOI:** 10.1101/2022.11.10.516058

**Authors:** Anupam Anand Ojha, Ambuj Srivastava, Lane William Votapka, Rommie E. Amaro

## Abstract

Janus kinases (JAK) are a group of proteins in the non-receptor tyrosine kinase (NRTKs) family that play a crucial role in growth, survival, and angiogenesis. They are activated by cytokines through the Janus kinase - signal transducer and activator of transcription (JAK-STAT) signaling pathway. JAK-STAT signaling pathways have significant roles in the regulation of cell division, apoptosis, and immunity. Identification of the V617F mutation in the Janus homology 2 (JH2) domain of JAK2 leading to myeloproliferative disorders has stimulated great interest in the drug discovery community to develop JAK2-specific inhibitors. However, such inhibitors should be selective towards JAK2 over other JAKs and display an extended residence time. Recently, novel JAK2/STAT5 axis inhibitors (N-(1H-pyrazol-3-yl)pyrimidin-2-amino derivatives) have displayed extended residence times (hours or longer) on target and adequate selectivity excluding JAK3. To facilitate a deeper understanding of the kinase-inhibitor interactions and advance the development of such inhibitors, we utilize a multiscale Markovian milestoning with Voronoi tessellations (MMVT) approach within the Simulation-Enabled Estimation of Kinetic Rates v.2 (SEEKR2) program to rank-order these inhibitors based on their kinetic properties and further explain the selectivity of JAK2 inhibitors over JAK3. Our approach investigates the kinetic and thermodynamic properties of JAK-inhibitor complexes in a user-friendly, fast, efficient, and accurate manner compared to other brute force and hybrid enhanced sampling approaches.

## 1 Introduction

Tyrosine kinases (TKs) are a family of proteins that catalyze the transfer of phosphate groups from adenosine triphosphate (ATP) molecules to tyrosine residues of the target protein.^1, 2^ The TKs can be broadly divided into receptor and non-receptor tyrosine kinases. The receptor tyrosine kinases (RTKs) are membrane-bound and pass the extracellular signal to the inside of cells, while non-receptor tyrosine kinases (nRTKs) are mainly cytosolic and bind to ligands to activate downstream signaling.^3–5^ nRTKs are involved in cell signaling, differentiation, proliferation, and apoptosis. Janus kinase (JAK) proteins are nRTK receptors involved in activating transcription and production of cytokines to recruit immune cells at the site of infections. The JAK family comprises of Janus kinase 1 (JAK1), Janus kinase 2 (JAK2), Janus kinase 3 (JAK3), and Tyrosine kinase 2 (TYK2).^6, 7^ JAKs regulate downstream signalling by activating signal transducer and activator of transcription (STAT) proteins propagating the signal from the membrane to the nucleus, also known as the JAK-STAT pathway.^7–9^ The JAK-STAT pathway regulates cytokines and growth hormones which are crucial for cellular processes, such as hematopoiesis, lactation, immune system development and immune response.^10^ The abnormalities and mutations in JAK proteins lead to neurological and immune system defects, including, but not limited to, rheumatoid arthritis (RA), inflammatory bowel diseases (IBD), multiple sclerosis (MS), and cancer.^11, 12^ Mutations in JAK1 and JAK3 are especially known to cause severe combined immune deficiency (SCID) diseases.^13^

The JAK proteins are constitutively expressed, with the exception of JAK3, which are expressed upon immune activation. JAK proteins contain seven conserved homology domains (JH1-JH7).^14, 15^ The JH1 domain at the carboxyl-terminal show a classical tyrosine kinase activity, while the JH2 domain are pseudo-kinase domains that assist the JH1 domain for catalysis. JH3-JH7 domains are known to be involved in receptor binding and the regulation of kinase activity. Inhibition of JAK proteins may prove to be effective against diseases, including neurological disorders and different types of cancer. The similarity and structural conservation in JAK proteins create challenges to designing selective inhibitors against them.^16, 17^ Although both the JAK2 and JAK3 proteins have highly conserved domains and are structurally very similar, one of the significant differences between them is the interaction of these proteins with different types of receptors. While JAK2 primarily mediates signals from glycoprotein 130 (gp130)-related cytokines, granulocyte macrophage-colony stimulating factor (GM-CSF) receptors, and type II cytokine receptors, JAK3 mediates signaling from type I receptors containing the common gamma chain (*γ*c).^18–21^ JAK inhibitors have shown promise as potential treatments for a variety of diseases, including certain types of cancer, autoimmune disorders, and inflammatory conditions.^22–27^ Tofacitinib and baricitinib are the two first-generation drugs that the federal drug administration (FDA), and the European medicines agency (EMA) have approved for the treatment of RA.^28–30^ Tofacitinib targets JAK1, JAK2, and JAK3, while baricitinib targets JAK1 and JAK2 proteins. However, selective inhibition of JAK proteins is crucial for tuning the signaling pathway and the underlying downstream processes. Structural understanding of selective inhibition is crucial to optimize their activities and design better selective inhibitors.^31^

Molecular dynamics (MD) simulations have been effective in studying the binding and unbinding dynamics of protein-inhibitor complexes and can be used for kinetic estimates.^32–43^ Understanding the receptor-ligand binding and unbinding process can be useful for drug discovery and development, especially in accelerating lead optimization efforts and lowering drug attrition rates.^44–46^ The bimolecular association rate constant (*k*_on_) and the dissociation rate constant (*k*_off_) are required to describe a kinetic profile of a potential noncovalent inhibitor or a drug molecule. Recently, drug-target residence time (1*/k*_off_), or the time spent by the drug in the binding pocket of the protein, has received significant attention as drugs with a higher residence time are shown to have greater *in vivo* efficacy as compared to thermodynamic parameters such as free energy. ^47–50^ It is possible for drugs with similar binding free energies (Δ*G_bind_*) to have different binding and unbinding kinetic rates. Several factors contribute to ligand binding and unbinding kinetics. These include, but are not limited to the size and flexibility of ligands, forces within the molecular system, large-scale receptor conformational rearrangements, and ligand-induced conformational changes in the receptor.^51–57^ One of the major limitations of MD simulations is the immense amount of computation required to observe rare biologically relevant events. Simulations often get stuck in metastable regions. Enhanced sampling methods, including and not limited to, metadynamics,^58–62^ adaptive biasing force (ABF),^63–65^ and umbrella sampling^66, 67^ are employed to overcome such limitations where the applied bias potential steers the system to overcome deep energy wells. The bias potential for these methods is a function of collective variables (CVs), which are predefined and often require an in-depth understanding of the biological systems of interest.

Gaussian accelerated molecular dynamics (GaMD) is an enhanced sampling method where a harmonic boost potential is added to the total potential energy of the system, leading to reduced energy barriers.^68, 69^ An implementation of GaMD for receptor-ligand complexes is Ligand GaMD (LiGaMD), where a potential energy boost is applied to the ligand non-bonded interaction potential energy while another boost is applied to the remaining potential energy of the entire system, thus facilitating accelerated ligand binding and unbinding events.^70, 71^ Random acceleration molecular dynamics (RAMD) is another method used to rank inhibitors by residence time for a particular receptor.^40, 72^ Scaled MD is an unbiased sampling approach that can be used to predict protein-ligand unbinding kinetics.^36^ Other methods, including free energy perturbation, can be used to obtain thermodynamic, but not kinetic, predictions for receptor-ligand binding.^73–76^ A number of enhanced sampling methods exist to predict the kinetics and thermodynamics of binding and unbinding and have been summarized in recent literature.^43, 77–82^ A study using MM-GBSA was recently performed on a similar kinase for inhibitors bound to the ATP binding site. ^83^ In contrast to biasing potential methods, for the JAK systems examined in this study, Simulation-Enabled Estimation of Kinetic Rates v.2 (SEEKR2) employs a reasonably simple and uniform CV definition for receptor-ligand complexes and requires a minimal *a priori* understanding of these complexes.

N-(1H-pyrazol-3-yl)pyrimidin-2-amino derivatives are analogous to ATP molecules and have been shown to selectively inhibit JAK2 proteins with a high residence time in the binding pocket of JAK2 as compared to JAK3 proteins (Figure 1).^84^ We, therefore, aim to rank these inhibitors in comparison to their experimentally reported residence times in the JAK complexes by employing a milestoning simulation method and explain the differences in residence times by providing a complete kinetic and thermodynamic profile of receptor-ligand pairs. The SEEKR2 software is user-friendly, fast, efficient, and accurate as compared to other brute force methods and hybrid approaches. ^68, 70, 85–90^ The Markovian milestoning with Voronoi tesselation (MMVT) method implemented in the SEEKR2 program is described in the Methods section followed by a detailed description of the calculation of residence times, and kinetic and thermodynamic profiles of the protein-inhibitor complexes.

**Figure 1:**
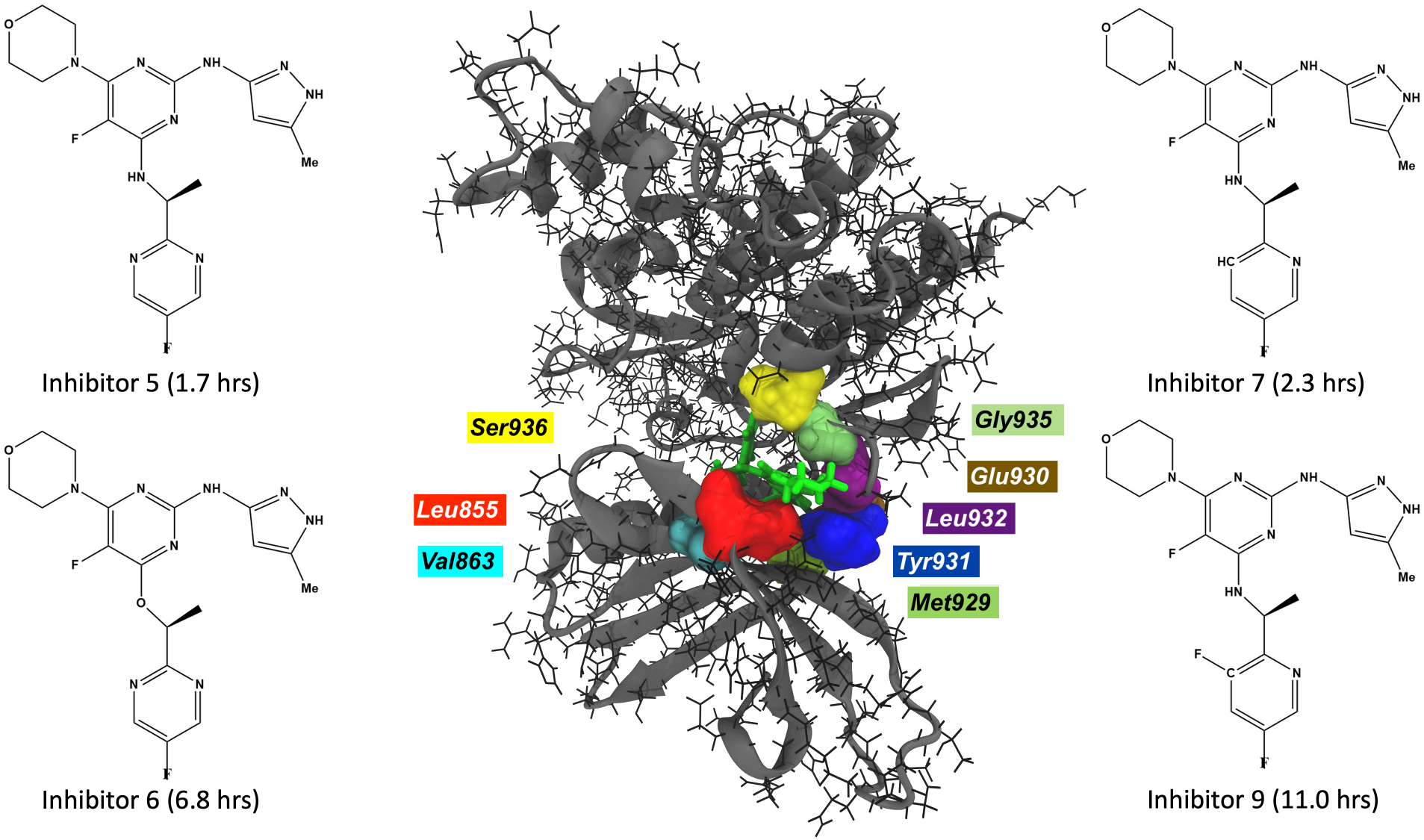
JAK2-inhibitor 9 complex with interacting residues within a cut-off distance of 2.5 Å (center). The inhibitors with large residence times for JAK2 proteins are displayed.

## 2 Methods

### 2.1 Simulation Enabled Estimation of Kinetic Rates v.2 (SEEKR2)

#### 2.1.1 Markovian milestoning with Voronoi tessellations

A Voronoi tessellation is a subdivision of space into n regions or “Voronoi cells”.^91, 92^ From a given set of points, **a** = *{a*_1_*, a*_2_*, a*_3_*, …., a_n_}* and set of Voronoi cells, **V** = *{V*_1_*, V*_2_*, V*_3_, …*, V_n_}* such that *a*_1_ *∈ V*_1_, *a*_2_ *∈ V*_2_, *a*_3_ *∈ V*_3_, …., *a_n_ ∈ V_n_* (Figure 2), let us define a distance metric, *d*(*a, b*), that estimates the distance between the two points, *a* and *b*. According to the definition of a Voronoi tessellation, a point *α* will belong to the cell *V*_1_ if and only if *d*(*a*_1_*, α*) *< d*(*a_i_*)*, α*) for *i ∈ {*2, 3*, …., n}*. There are *N* boundaries (milestones) between adjacent Voronoi cells.

**Figure 2:**
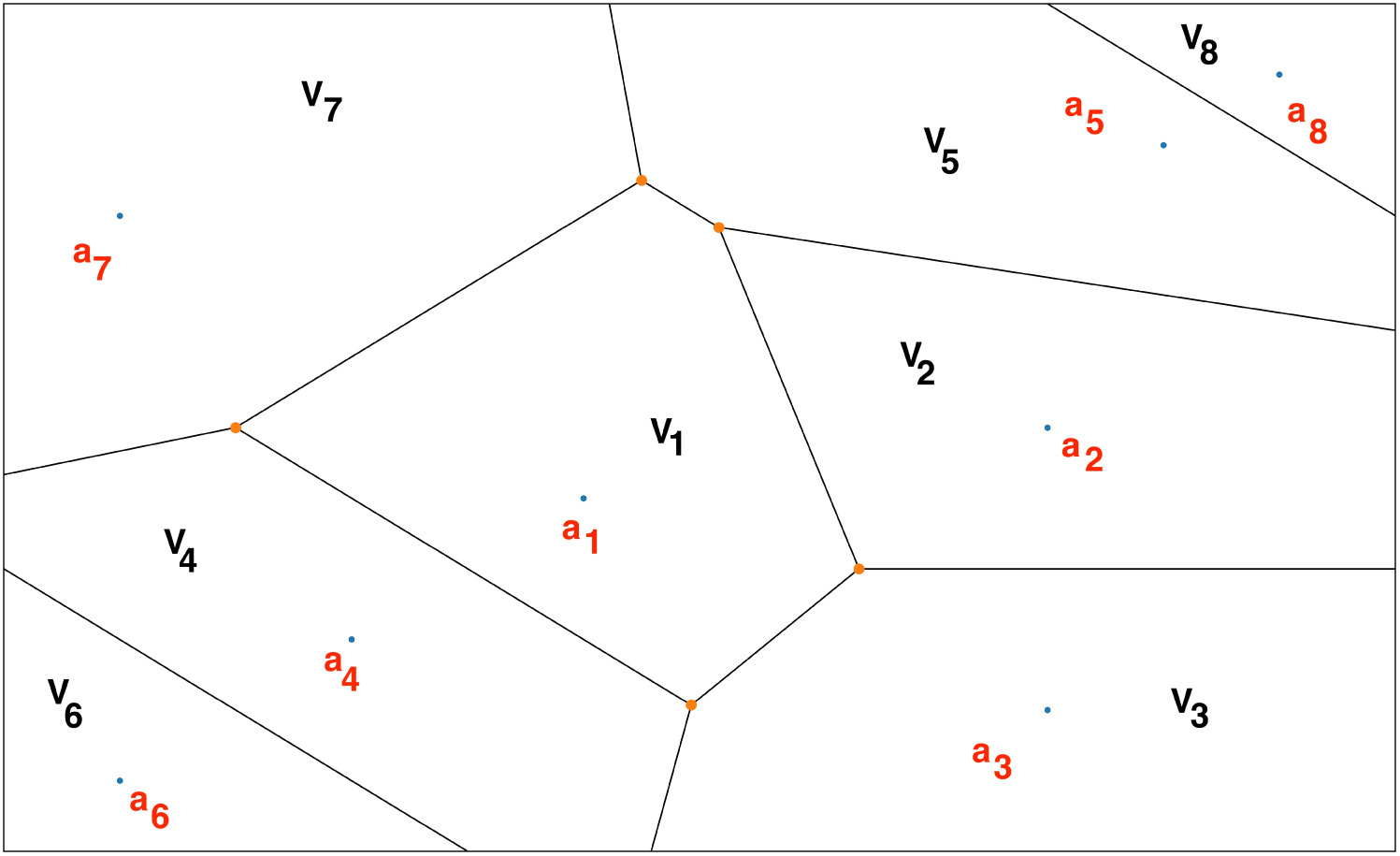
A representative Voronoi diagram where *V*_1_*, V*_2_*, V*_3_*, …., V_n_* represents the Voronoi cells and *a*_1_ *∈ V*_1_, *a*_2_ *∈ V*_2_, *a*_3_ *∈ V*_3_,*….*, *a_n_∈ V_n_*.

SEEKR2 is an open-source software that automates the process of preparation, initiation, running, and analysis of milestoning calculations utilizing MD and Brownian dynamics (BD) simulations to estimate the kinetics and thermodynamics of receptor-ligand binding and unbinding.^93–95^ MD simulations are run using the OpenMM simulation engine, while BD simulations are run using the Browndye software. ^96^ In the SEEKR2 multiscale milestoning approach, the phase space of the receptor-ligand complex is split into two regions, i.e., the MD and the BD region. This partition is based on a predefined CV, i.e., the distance between the center of mass (COM) of the ligand and the COM of the receptor’s binding site. In the region closer to the binding site, solvent effects and molecular flexibility must be included for describing molecular interactions, therefore, MD simulations are employed. The MD region is further partitioned into several Voronoi cells. Steered molecular dynamics (SMD) simulations are run to generate starting structures for SEEKR2 simulations. ^97^ SMD simulations pull the ligand slowly out of the binding pocket with a moving harmonic restraint, and a snapshot of the trajectory is saved for every Voronoi cell as it passes through them. Fully atomistic, flexible, and parallel MD simulations are performed in each Voronoi cell with reflective boundary conditions. When the ligand is further away from the binding site, i.e., in the BD region, rigid body BD simulations are adequate to describe the diffusional encounter of the ligand and the receptor.

The MMVT-SEEKR2 approach has been shown to estimate binding and unbinding kinetic and thermodynamic properties for less complex receptor-ligand systems with close accuracy, especially the model host-guest systems, i.e., *β*-cyclodextrin with guest ligands and the model protein system, i.e., the trypsin-benzamidine complex.^95^ We thereby extend our efforts in exploring the capabilities of SEEKR2 in estimating kinetic and thermodynamic properties for more complex systems, specifically ligands which are strong binders and have large residence times.

#### 2.1.2 Estimating residence times and binding free energies

According to the MMVT approach, the system evolves according to a continuous-time Markov jump between Voronoi cells.^98, 99^ Let the rate matrix associated with the evolution be **Q**, *N* _ij_ be the number of transitions between milestones, *i* and *j*, and *R*_i_ be the time spent by the trajectory having last touched milestone *i*. The diagonal and the off-diagonal elements of the transition matrix, **Q** are represented by equations 1 and 2 respectively.

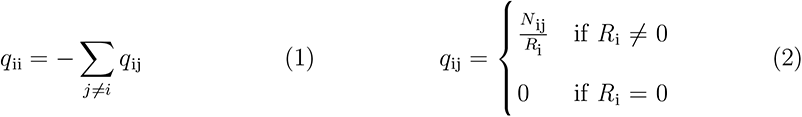

MD simulations are run within the Voronoi cells until convergence is reached. Reflective boundary conditions are employed at the boundaries to confine trajectories within the Voronoi cells. Consequently, velocities of the trajectories are reversed as they touch the edges of the adjacent Voronoi cells. For a Voronoi cell *α*, let *N^α^* be the number of trajectory collisions with an *i* ^th^ milestone after having last touched the *j* ^th^ milestone within anchor *α*, let *R^α^* be the simulation time having last touched the *i* _th_ milestone within anchor *α*, let *T_α_* be the total simulation time in cell *α*, let *N_α,β_* be the total number of collisions within Voronoi cell *α*, with the boundary shared with Voronoi cell *β*, and let *T* be the reciprocal sum of time spent in all the cells as described by equation 3, then *N* _ij_ and *R*_i_ is represented by equations 4 and 5 respectively. The equilibrium probability, *π* is obtained by solving equations 6 and 7.

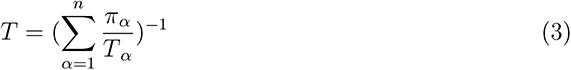

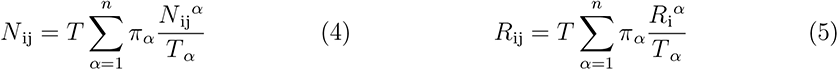

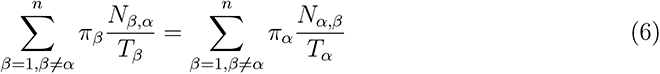

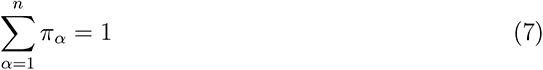

With **Q** as the *N* -1 by *N* -1 matrix obtained from the upper left corner of **Q**, one can compute the mean first passage time (MFPT) or residence time for each milestone described by vector **T***^N^* by solving equation 8.

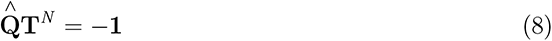

where **1** is a vector of ones. Stationary probabilities obtained from the milestoning simulations are used to construct the free energy profile of unbinding of the receptor-inhibitor complexes with the bound-state milestone as a reference. Stationary probabilities, **p**, are found by solving the eigenvalue equation 9.

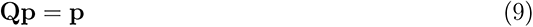

Let k_B_ be Boltzmann’s constant, *T* be the temperature, *p_i_* be the stationary probability of the i_th_ milestone and *p_ref_* be the stationary probability of the bound-state or the reference milestone. The expression for estimating the free energy profile of the i^th^ milestone, i.e., Δ*Gi*_I_ is given by equation 10.

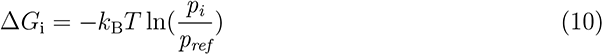

#### 2.1.3 Ranking JAK-inhibitor complexes with SEEKR2

N-(1H-pyrazol-3-yl)pyrimidin-2-amino derivatives are ATP-competitive inhibitors of the JAK2-STAT5 pathway that are reported to display prolonged residence times on JAK2 and sufficient selectivity against JAK3, both at biochemical and cellular levels. ^84, 100^ The residence times of four inhibitors with the JAK2 and JAK3 kinase, were experimentally determined using a rapid dilution enzymatic assay. ^101^ We present a relatively computationally inexpensive and efficient application of the SEEKR2 program to predict and rank order the residence times of the JAK2 and JAK3 inhibitors.

##### System Preparation

To estimate the residence times of the four inhibitors in the two JAK proteins, an all-atomistic MD simulation is performed in SEEKR2. The X-ray crystal structure of JAK2-JH1 domain in complex with inhibitor 6 (PDB ID: 3ZMM) is used as the reference structure for JAK2 SEEKR2 simulations.^84^ For the preparation of JAK2 complex with inhibitors 5, 7, and 9 (Figure 1), the X-ray crystal structure of JAK2 domain in complex with inhibitor 6 (PDB ID: 3ZMM) is used as a reference structure. Inhibitor 6 is modified to 5, 7, and 9 using the Maestro module of the Schrödinger software suite (Figure 1). ^102^ Once inhibitor 6 is modified to either inhibitor 5, 7, or 9, the JAK-inhibitor complex is subjected to the removal of water molecules beyond 3 Å of the protein and with fewer than three hydrogen bonds to the neighboring residues. It is followed by hydrogen bond optimization of the receptor-ligand complex with protonation states of residues at pH=7.4. Finally, a restrained minimization of the complex is performed with a complete relaxation of the H-bond network while keeping the heavy atoms restrained. The AMBER ff14SB forcefield is used to parameterize the protein, while the inhibitor is parameterized using the Antechamber module with the general Amber force field (GAFF) with the AM1-BCC charge model.^103–106^ The protein-inhibitor complex is then explicitly solvated with the TIP4P-Ew water model and a salt (Na+/Cl-) concentration of 150 mM in a truncated octahedral periodic box with a 10 Å water buffer.^107^ The OpenMM MD engine is used to run the simulation at 300 K with a 2 fs timestep and a nonbonded cut-off radius of 9 Å.^108, 109^ The system is systematically heated from 0 K to 300 K in steps of 3 K of 20 ps each, followed by 20 ns each of NPT and NVT equilibration simulations.

The X-ray crystal structure of JAK3-JH1 domain in complex with an indazole substituted pyrrolopyrazine (PDB ID: 3ZC6) is used as the reference structure for the JAK3 SEEKR2 simulations.^110^ The inhibitor complexed with JAK3 is removed, and the structure is aligned to the JAK2 complexed with inhibitor 6. Inhibitor 6 is then placed at the ATP binding site of the JAK3 protein. Inhibitor 6 is modified to 5, 7, and 9 using the Maestro module of the Schrödinger software suite, and the same protocol is followed for JAK3 systems as performed for the JAK2 complexes for system preparation, solvation and equilibration. It is important to note that only one crystal structure of JAK2 is used to prepare all four JAK2-inhibitor complexes, and the same holds true for the JAK3-inhibitor complexes.

##### Steered molecular dynamics and Voronoi cell definition

To define Voronoi cells, we described the CV as the distance between the COM of the inhibitor and the COM of *α*-carbons of the binding site^111^ (Supplementary Table S1). The cut-off distance for the binding site of the inhibitor is defined as 3 Å. All the *α*-carbon atoms of the surrounding residues of the JAK protein within the cut-off distance of any of the atoms of the inhibitor is defined as the binding site for the receptor-inhibitor complex. Supplementary Table S1 displays the residues of each JAK-inhibitor complex selected for the COM calculation of the binding site. For JAK2-inhibitor complexes, CV-based milestones are defined as concentric spheres and are located at distances of 2.5, 3.0, 3.5, 4.0, 4.5, 5.0, 5.5, 6.0, 6.5, 7.0, 7.5, 8.0, 8.5, 9.0, 9.5, 10.0, 11.0, 12.0, 13.0, 14.0, 15.0, and 16.0 Å respectively from the COM of the binding site. Similarly, for the JAK3-inhibitor complexes, CV-based milestones are defined as concentric spheres and are located at a distance of 3.0, 3.5, 4.0, 4.5, 5.0, 5.5, 6.0, 6.5, 7.0, 7.5, 8.0, 8.5, 9.0, 9.5, 10.0, 11.0, 12.0, 13.0, 14.0, 15.0, and 16.0 Å respectively from the COM of the binding site. In the case of JAK3-inhibitor complexes, none of the residues of the JAK3 protein interacted with the inhibitor within the 2.5 Å radius, leading to the choice of the first milestone at 3.0 Å. This choice should not be problematic since the milestoning procedure would not be significantly sensitive to the choice of the number of milestones as long as each state and pathway is adequately represented in each milestoning model and the results are sufficiently converged. SMD simulations are employed to generate starting structures within each Voronoi cell where the ligand bound to the complex is slowly pulled out of the binding site in such a way that there is no significant stress to the system and it stays in the local equilibrium. To generate starting structures for MMVT simulations, the ligand is slowly pulled from the bound state to the outermost Voronoi cell with a moving harmonic restraint of 50,000 kJ mol^-1^ nm^-2^ over the course of 1 *µ*s.

##### SEEKR2 molecular dynamics simulations

With the starting structures of each Voronoi cell obtained by SMD, MMVT simulations are employed with the same forcefield parameter files used during equilibration simulations. No harmonic restraint is applied during these simulations. Reflective boundary conditions are employed to retain the trajectories within individual Voronoi cells. A total of 400 ns of MD simulations is run within each Voronoi cell. To improve the sampling and account for stochasticity, three replicas of SEEKR2 simulations are run for each JAK-inhibitor complex. In short, three replicas of 21 independent and parallel MD simulations of 400 ns are run for each of the JAK2-inhibitor complexes, totaling a simulation time of 25.2 µs. Similarly, three replicas of 20 independent and parallel MD simulations of 400 ns within each Voronoi cell are run for each of the JAK3-inhibitor complexes, totaling a simulation time of 24 µs. For the JAK2-inhibitor and JAK3-inhibitor complexes, 21 and 20 parallel simulations, respectively for 400 ns each, were carried out on one NVIDIA V100 GPU on the Popeye computing cluster at San Diego Supercomputer Center (SDSC), which aggregated approximately 220 ns/day, i.e., the entire SEEKR2 simulations for each complex required approximately 44 hrs. of computing on parallel GPUs (21 and 20 parallel GPUs for JAK2-inhibitor and JAK3-inhibitor complexes respectively). Therefore, SEEKR2 is a powerful tool for rank ordering the ligands and characterizing the ligand binding and unbinding kinetics and thermodynamics in receptor-ligand complexes in a user-friendly and computationally efficient manner, thus facilitating computer-aided drug design.

## 3 Results and Discussion

Estimating thermodynamic and kinetic parameters, such as the residence time and free energy of binding and unbinding, is challenging in cases of receptor-inhibitor complexes with an extended residence time.^112, 113^ A minor change in the structure of inhibitors sometimes leads to an enormous change in the residence time in the binding pockets of proteins. We estimated the residence time of four inhibitors in the binding pocket of JAK2 and JAK3 proteins. We showed that the trend of the residence time predicted by the SEEKR2 milestoning approach captures that of experimental methods. We showed that the trend of the residence time predicted by the SEEKR2 milestoning approach reproduces the experimental findings. Inhibitors 5 and 9 displayed the lowest and the highest residence times for the JAK2 protein, respectively. Similarly, inhibitors 6 and 9 displayed the lowest and the highest residence times for the JAK3 protein, respectively (Supplementary Table S2). Long timescale MD simulations are performed to study the structural aspects of protein-ligand interactions, primarily focusing on these particular inhibitors to explain the discrepancy in their respective residence times.

### 3.1 Determination of kinetic and thermodynamic parameters from SEEKR2 simulations

Simulations in the majority of the Voronoi cells converged after 400 ns. The MFPT or residence time is calculated using equation 8. The residence times reported in Figure 3a and 3b are the means of the residence times obtained from three independent SEEKR2 simulations for each of the JAK-inhibitor complexes (Supplementary Table S2). Residence times for the novel series of inhibitors for the JAK2 and JAK3 estimated by the SEEKR2 program are in close agreement with the experimental studies (Figure 3a and 3b). SEEKR2 not only predicted the residence times correctly but also preserved the rank ordering of residence times for inhibitors in both the JAK2 and JAK3 complexes. It can be seen from Figure 3a and 3b that inhibitors 6 and 9 display extended residence times in the ATP-binding site of the JAK2 complexes.

**Figure 3:**
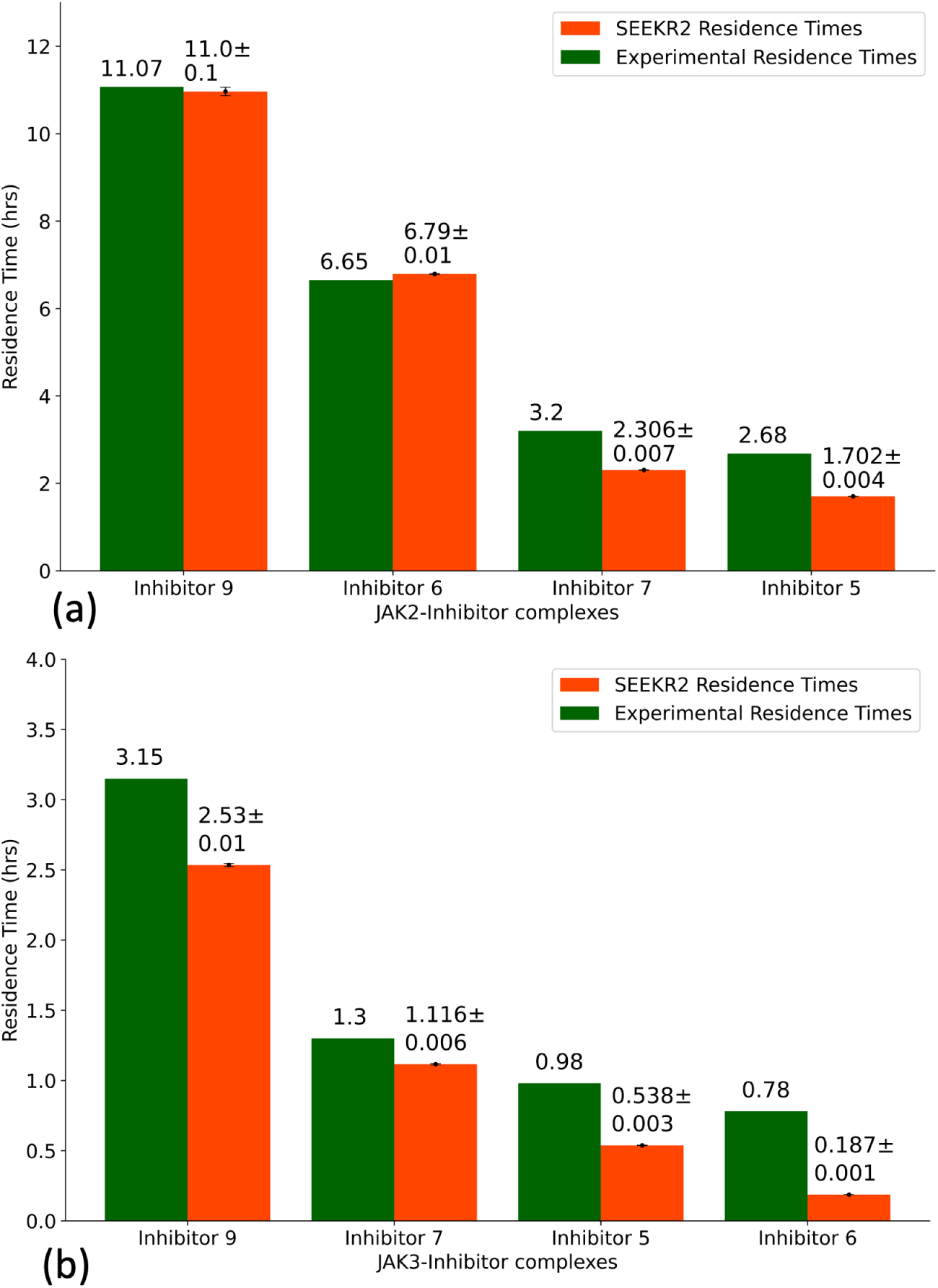
Residence times of JAK2 and JAK3 inhibitors as obtained from the experiments and the SEEKR2 milestoning method. The values of the residence times and the error bars for each JAK-inhibitor complex is the average of the three independent SEEKR2 calculations. (a) Residence times of the inhibitors for the JAK2 protein and (b) residence times of the inhibitors for the JAK3 protein are displayed in the figure. Error bars are present for the SEEKR2 residence time data, but they are sometimes too small be visible in the figure. An unpaired t-test is carried out to measure the statistical significance of the difference between the experimentally determined residence times of JAK2 and JAK3 inhibitors and the SEEKR2-calculated residence times. The p-values obtained from the t-test determined that there is no significant difference between the mean of the SEEKR2-calculated residence times and the experimentally determined residence times (Supplementary Table S3).

ΔG_i_ is calculated for each of the milestones using equation 10. In the case of the JAK2-inhibitor 5 complex, two energy barriers exist as the inhibitor dissociates with the receptor, one at milestone 4 and the other at milestone 11 (Figure 4a). The COM-COM distance between the inhibitor and the alpha-carbon (*α*-C) atoms of the binding site for the first transition state (TS 1) is 4.50 Å, while the second transition state (TS 2) is at a COM-COM distance of 8.00 Å from the binding site. Similarly, two energy barriers exist for the JAK2-inhibitor 9 complex, one at milestone 5 and the other at milestone 13 (Figure 4a). TS 1 is at a COM-COM distance of 5.00 Å, while TS 2 is at a COM-COM distance of 9.00 Å from the binding site. The energy barriers for inhibitor 9 for both transitions are higher than that of inhibitor 5, indicating that inhibitor 9 is a stronger binder with a higher residence time. For the JAK3-inhibitor 6 complex, two energy barriers exist as the inhibitor dissociates with the receptor, one at milestone 7 and the other at milestone 12 (Figure 4b). The COM-COM distance between the inhibitor and the *α*-C atoms of the binding site for TS 1 is 6.50 Å, while TS 2 is at a COM-COM distance of 9.00 Å. Similarly, two energy barriers exist for the JAK3-inhibitor 9 complex, one at milestone 5 and the other at milestone 11 (Figure 4b). TS 1 is at a COM-COM distance of 5.00 Å, while TS 2 is at a COM-COM distance of 8.50 Å from the binding site. The energy barrier for inhibitor 9 for TS 1 is higher than that of inhibitor 6, indicating that inhibitor 9 is a stronger binder with a higher residence time.

**Figure 4:**
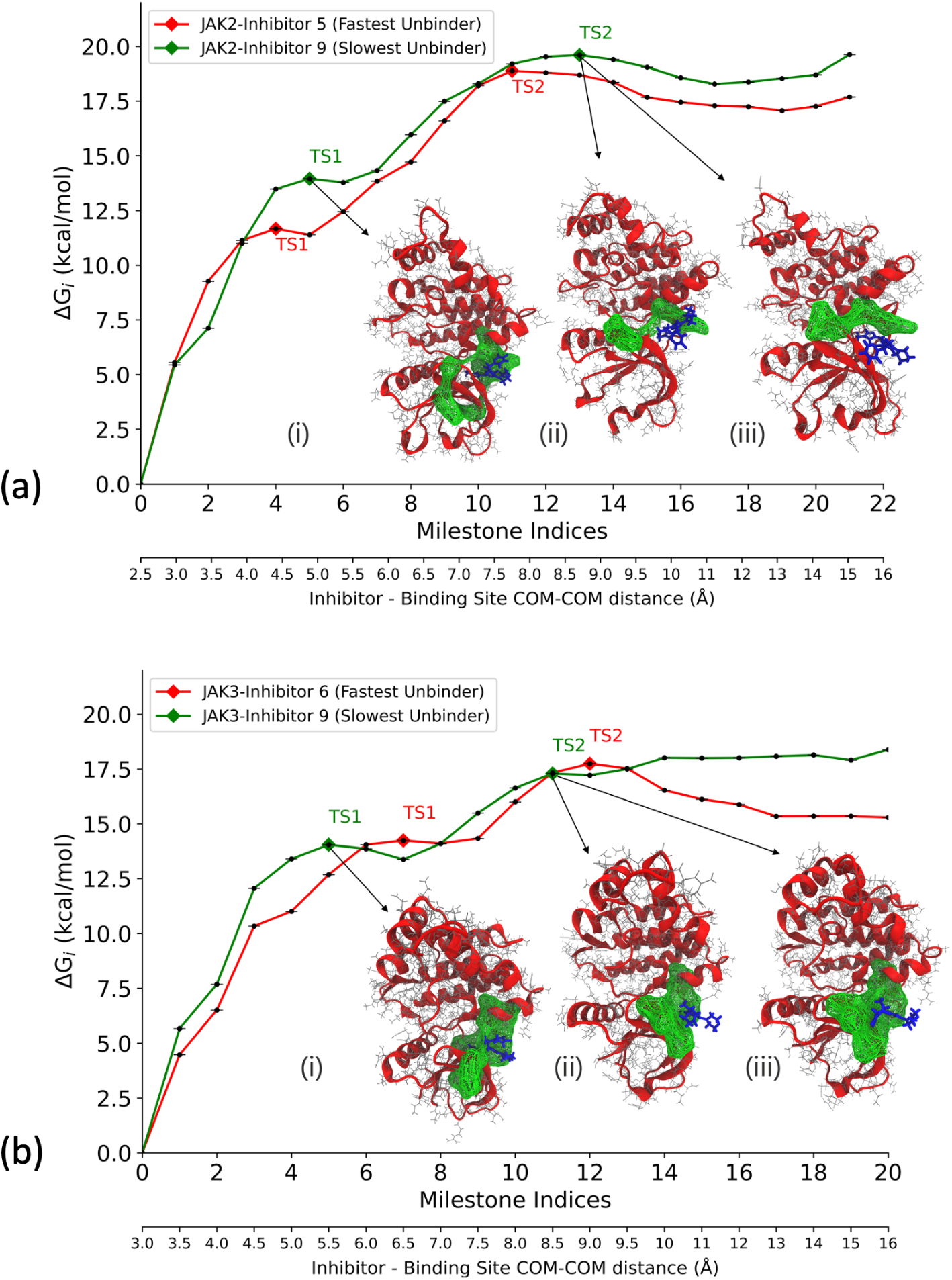
The free energy profile (ΔG_i_) obtained from the SEEKR2 milestoning method for the JAK proteins complexed with the inhibitors. Also shown are the dominant poses of inhibitor 9 as it unbinds from the ATP binding site of JAK complexes. These poses are obtained from the SEEKR2 trajectories for milestones with the local maximum values of ΔG_i_. ΔG_i_ values obtained for each of the JAK-inhibitor complex is the average of the three independent SEEKR2 calculations. The additional X-axis at the bottom of the graph denotes the distance between the center of masses of the inhibitor and the alpha carbon atoms of the binding site for each milestone. (a) ΔG_i_ values for the JAK2 protein complexed with inhibitor 5 and inhibitor 9 along with (i) JAK2-inhibitor 9 complex at TS 1 (ii) JAK2-inhibitor 9 complex at TS 2 (pose 1) (iii) JAK2-inhibitor 9 complex at TS 2 (pose 2) (b) ΔG_i_ values for the JAK3 protein complexed with inhibitor 6 and inhibitor 9 along with (i) JAK3-inhibitor 9 complex at TS 1 (ii) JAK3-inhibitor 9 complex at TS 2 (pose 1) (iii) JAK3-inhibitor 9 complex at TS 2 (pose 2)

With SEEKR2 simulations, we hold the advantage of predicting a possible ligand unbinding pathway since this methodology enables the receptor-ligand complex to undergo parallel simulations with the ligand at increasing distances from the binding site. MD trajectories within the milestones located at these transition barriers are analyzed to identify important ligand-residue interactions. For the JAK2-inhibitor 9 and JAK3-inhibitor 9 complexes, hydrogen bond (H-bond) analysis is conducted for the two identified transition states using the CPPTRAJ module of the Amber 22 package.^114–116^ In the case of the JAK2-inhibitor 9 complex, for TS 1, Gly935, Tyr931, and Asp939 interacted significantly with inhibitor 9 as H-bond acceptors, while Ser936, Leu932, and Tyr931 residues were H-bond donors to inhibitor 9 (Figure 5a and 6a). On the contrary, for TS 2, interactions between the residues and inhibitor 9 decreased significantly, where the residues closer to the terminals interacted as the inhibitor gradually unbinds from the binding site (Figure 5b and 6b). In the case of JAK3-inhibitor 9 complex, for TS 1, Tyr904 and Leu905 interacted with inhibitor 9 as H-bond acceptors and donors simultaneously (Figure 5c). For TS 2, interactions between the residues and inhibitor 9 were still significant, including Leu828 and Gly908 as major donor residues (Figure 5d). Interestingly, more residues were involved in the H-bond interactions at TS1 for the JAK2-inhibitor 9 complex compared to the JAK3-inhibitor 9 complex. This observation can be attributed to the selectivity of inhibitor 9 towards the JAK2 protein.

**Figure 5:**
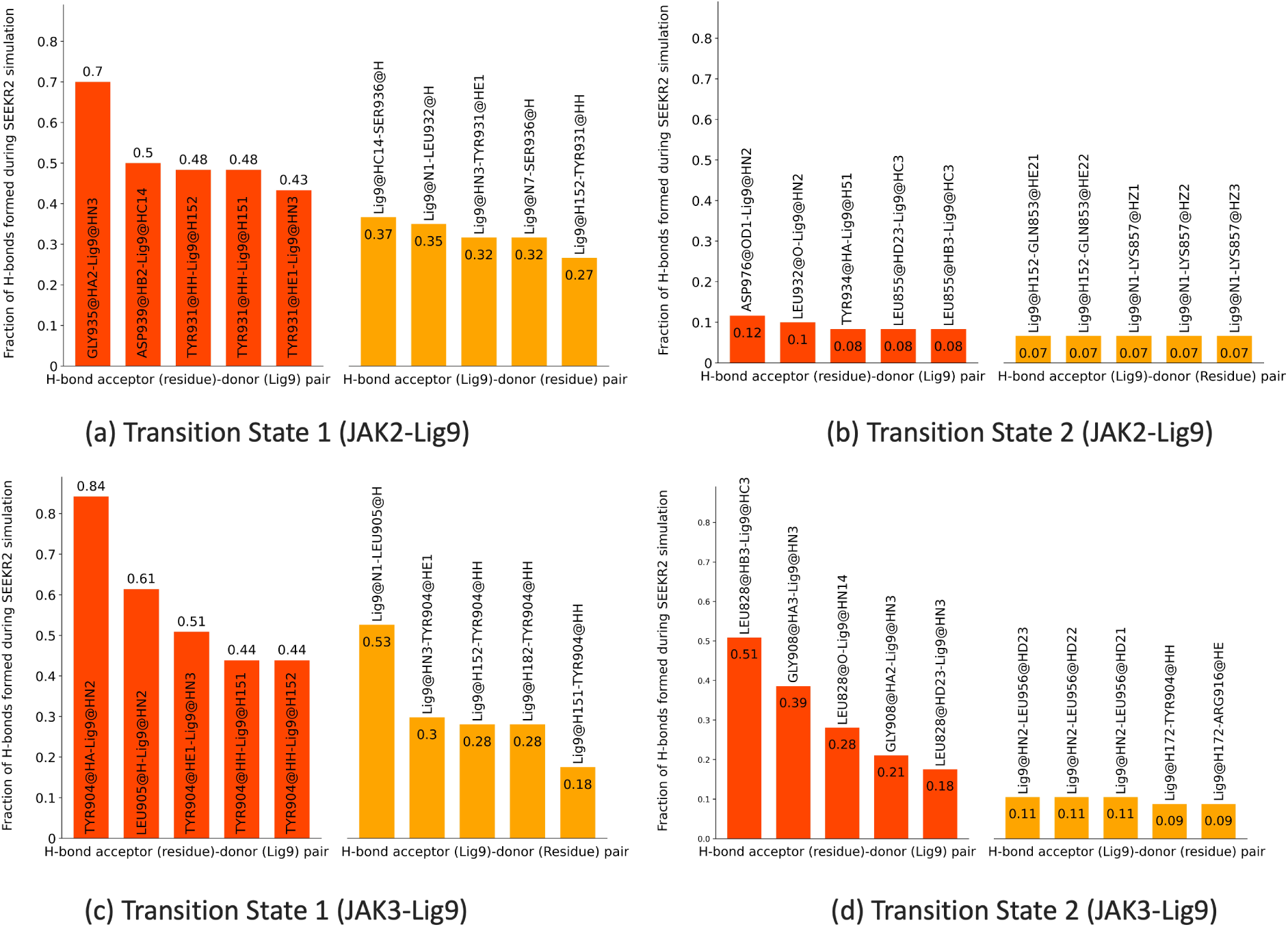
(a-b) Major hydrogen bond interactions formed during SEEKR2 simulations at transition states for the JAK2-inhibitor 9 complex displaying (a) TS 1 H-bond donor-acceptor pairs (b) TS 2 H-bond donor-acceptor pairs. (c-d) Major hydrogen bond interactions formed during SEEKR2 simulations at transition states for the JAK3-inhibitor 9 complex displaying (c) TS 1 H-bond donor-acceptor pairs (d) TS 2 H-bond donor-acceptor pairs.

**Figure 6:**
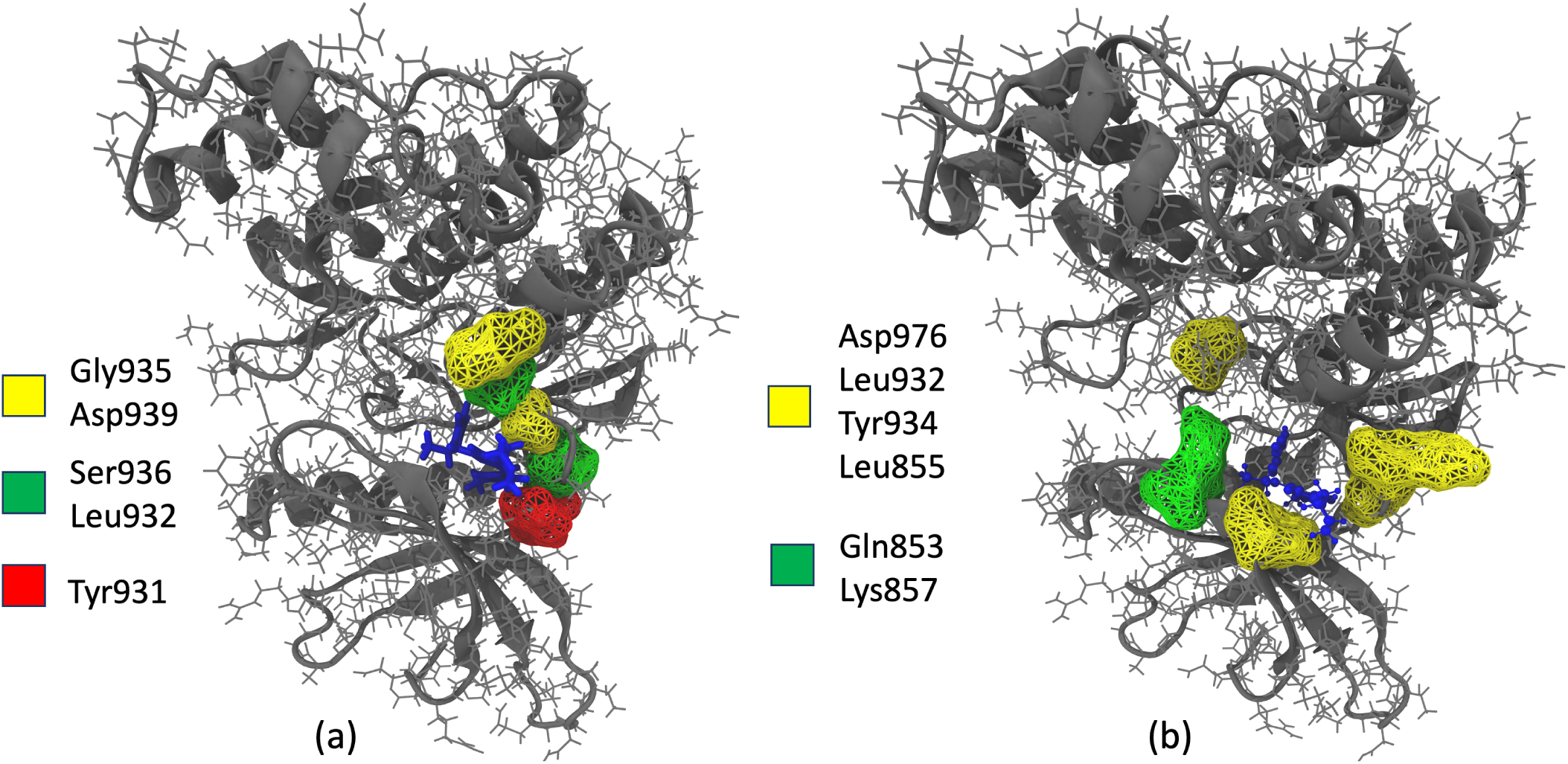
Major hydrogen bond interactions formed during SEEKR2 simulations for the JAK2-inhibitor 9 complex at (a) TS 1 displaying H-bond acceptor residues (yellow), H-bond donor residues (green), and H-bond donor/acceptor residues (red) (b) TS 2 displaying H-bond acceptor residues (yellow) and H-bond donor residues (green).

SEEKR2 is able to provide kinetic and thermodynamic estimates of receptor-ligand binding and unbinding, such as residence time and free energy of binding. Selectivity of inhibitors towards JAK2/JAK3 is a complex and multifaceted concept that cannot be reduced to a single physical quantity like residence time or free energy. Instead, it encapsulates the desirable outcome that the inhibitor more preferentially binds one potential target over another, which is influenced by numerous factors, including structural differences, conformational changes, off-target effects, cellular context, and pharmacokinetics.^117–120^ In this study, we focus on kinetic selectivity showing that SEEKR2 can discern a significant difference in residence times for the same set of inhibitors in JAK2 and JAK3. Recent literature studies show that thermodynamic and kinetic selectivity plays the most important role for targets of differing vulnerability, i.e., targets that require certain amounts of engagement with an inhibitor for the desired effect to be observed.^120–122^ Whether a target is high- or low-vulnerability depends, of course, on the desired effect. The actual mechanism of that selectivity is beyond the scope of the current study. Unfortunately, SEEKR2 alone is not able to discern the selectivity mechanisms, and additional analyses must be performed, as were performed in this study with the principal component analysis (PCA) and quantum mechanical calculations.

### 3.2 Long scale Molecular Dynamics Simulations

To understand and analyze critical aspects of binding and unbinding of the inhibitors at the ATP binding sites of JAK2 and JAK3 and to explain the discrepancy in the residence times of inhibitors and selectivity towards JAK2 over JAK3, three independent 2 *µ*s MD simulations are run for each JAK-inhibitor complex. The starting structures in the first Voronoi cell for each receptor-inhibitor complex served as the starting structures for the long-timescale MD simulations. We used the same forcefield parameter files for the complexes as used in the SEEKR2 simulations. For each of the receptor-inhibitor complexes, a total of 6 *µ*s of MD simulations are run at 300 K with a 2 fs timestep and a nonbonded cut-off radius of 9 Å using the OpenMM MD engine. Simulation trajectories are analyzed using the CPPTRAJ module of the Amber 22 package.^114–116^ Analyses including and not limited to ligand-binding site distance analysis, minimum average distance analysis, principal component analysis (PCA), and root mean squared fluctuation (RMSF) analysis are performed to gain a deeper understanding of the binding behavior of these inhibitors.

#### 3.2.1 Discrepancy in residence times: Structure of inhibitors and their interactions with JAKs

The inhibitors, namely 5, 6, 7, and 9, constitute a pyrazol-3-yl amine ring, a heteroaryl Cring, and a morpholine ring (Figure 7a). Different inhibitors are synthesized by substitutions at the heteroaryl C-ring. The pyrazol-3-yl amine ring forms multiple hydrogen bonds with the ATP binding pocket of the JAKs (Figure 8a and 8b), and these contacts are consistent with all the inhibitors. The solvent-exposed morpholine ring does not interact much with the residues in the binding region. Interestingly, a single substitution at the heteroaryl C-ring of the inhibitor leads to a significant difference in their residence times (Figure 8c). In the case of inhibitor 9 with respect to inhibitor 5, one of the nitrogen atoms in the heteroaryl C-ring is substituted by a -CF group (Figure 8c), leading to a five-fold increase in the residence time of inhibitor 9.

**Figure 7:**
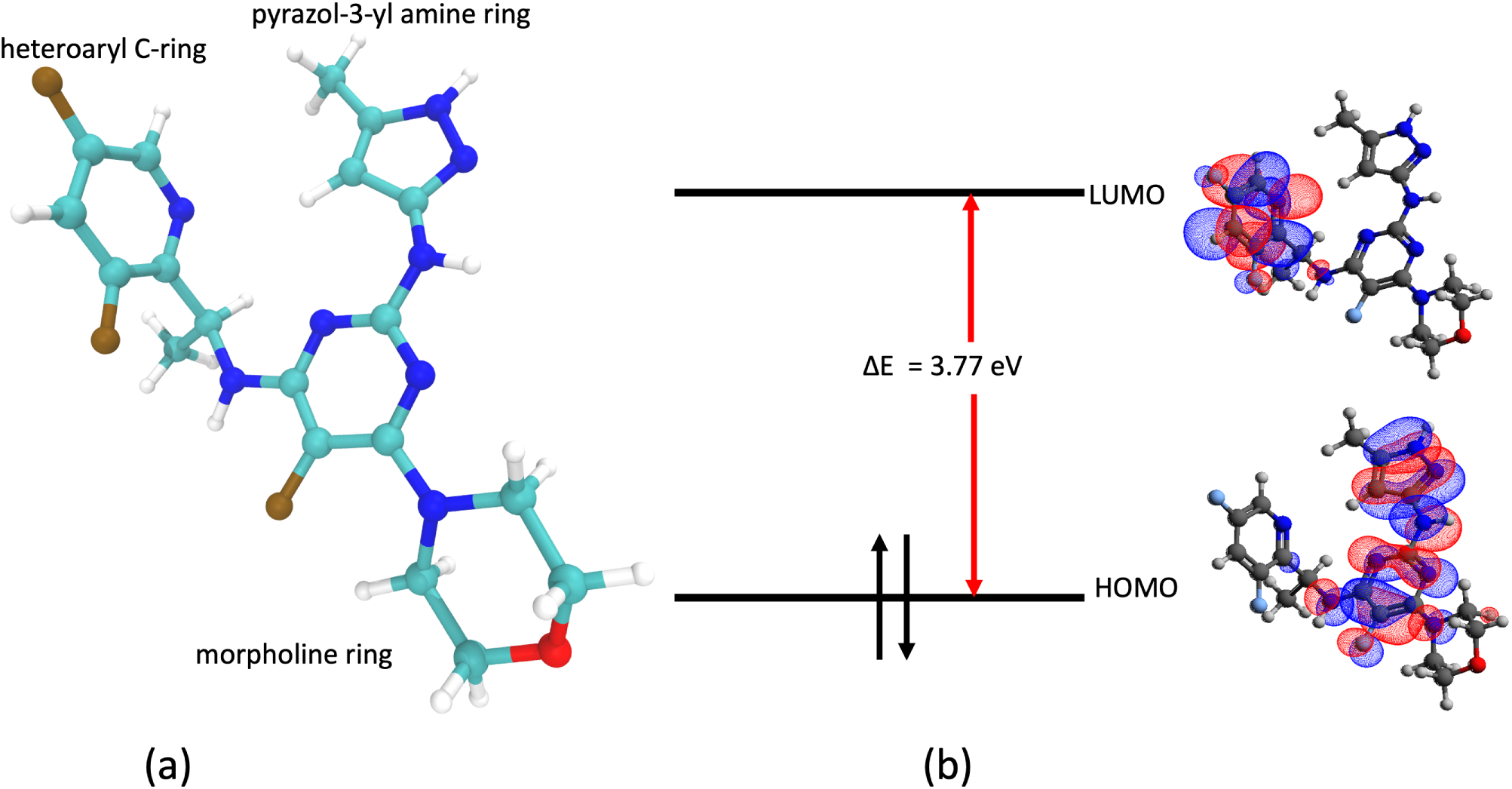
(a) Composition of inhibitor 9 and (b) Molecular orbitals of inhibitor 9

**Figure 8:**
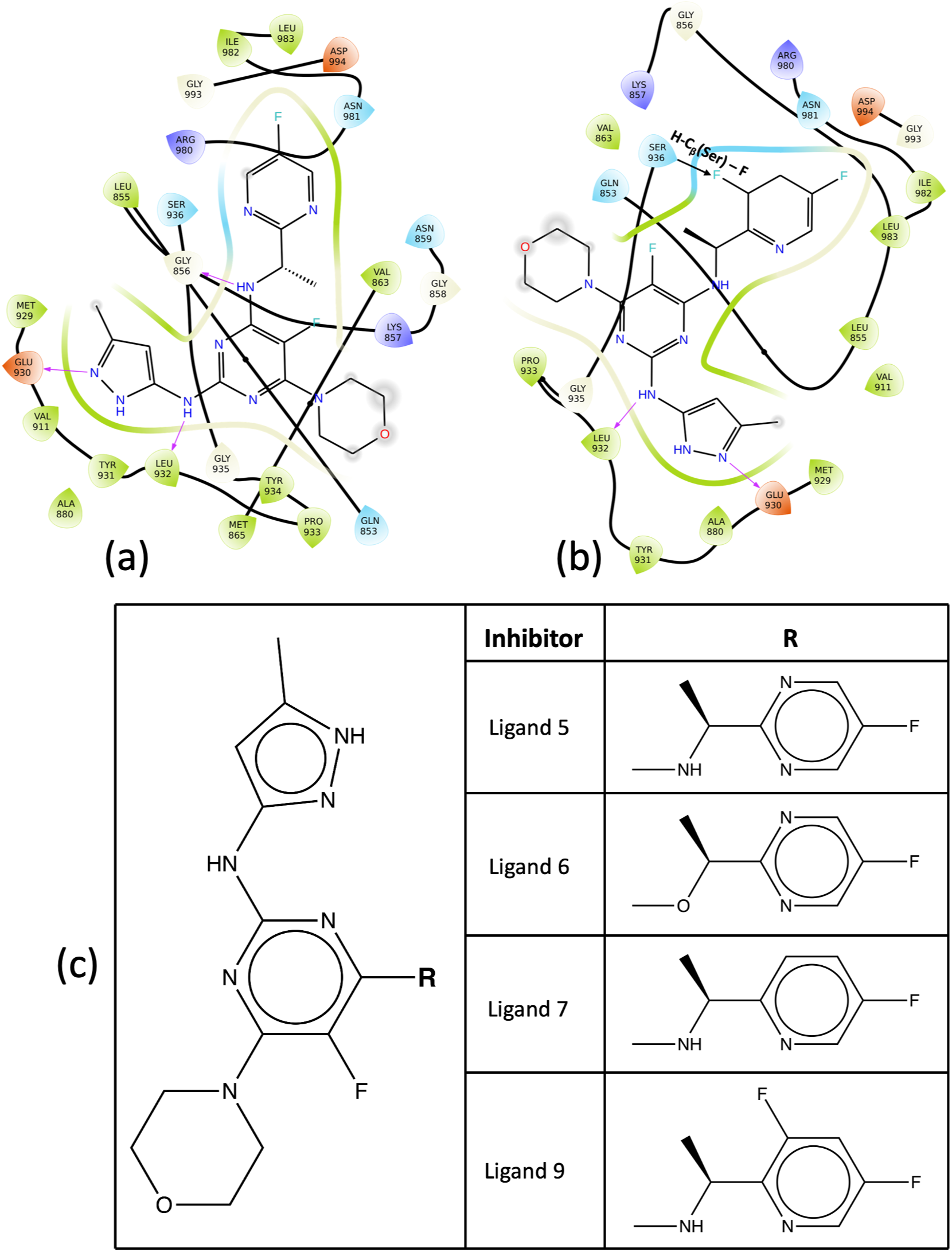
(a-b) Binding site of inhibitors for JAK2 complex showing important interactions with surrounding residues. (a) JAK2-inhibitor 5 complex (b) JAK2-inhibitor 9 complex (c) 2D formulae schemes for the JAK inhibitors indicating the location of modifications. In inhibitor 9, the substituted fluorine atom in the heteroaryl C-ring leads to the electrostatic pull of the hydrogen atom in the nearby serine residue, which contributes to the higher residence time in the kinase domain.

Inhibitor 9 displayed the highest residence time in both the JAK2 and JAK3 proteins. To investigate further the contributions of the heteroaryl C-ring towards the increased residence time and determine the donor-acceptor capabilities of the inhibitor, quantum mechanical (QM) calculations are run for inhibitor 5 and inhibitor 9 to determine the highest occupied molecular orbitals (HOMO) and lowest unoccupied molecular orbitals (LUMO). The Gaussian 16 suite of programs is used to carry out geometry optimization using Becke’s three-parameter functional in combination with the Lee-Yang-Parr correlation functional (B3LYP) and 6-31G(d,p) basis set.^123–126^ It is observed that the heteroaryl C-ring constitutes the LUMO (Figure 7b) for all the inhibitors. The presence of an extra fluorine atom in inhibitor 9 causes extra stabilization of the bound state since the substituted fluorine atom in the heteroaryl C-ring interacts with the hydrogen of the *β*-carbon of the serine residue (Ser936), maintaining an average distance of 2.64 Å with a minimum distance of 2 Å (Figure 8b). In contrast, for inhibitor 5, this interaction is missing (Figure 8a). Further evidence is provided by the HOMO-LUMO energy calculations obtained from the QM calculations. It is observed that the HOMO-LUMO energy difference for inhibitor 9 (3.77 eV) is higher than that of inhibitor 5 (3.40 eV). The HUMO energies for inhibitors 5 and 9 are nearly identical, but the LUMO energy for inhibitor 9 is higher than that for inhibitor 5. A higher energy LUMO suggests a more electron-deficient character of the heteroaryl C-ring leading to stabilization interaction with the serine (Ser936) residue of JAK2. In short, the electronegativity of F leads to the electrostatic pull of the hydrogen atom in the serine residue and is responsible for a higher residence time for inhibitor 9 than other inhibitors.

To gain additional insights into the dynamics of the receptor-inhibitor complex and to explain the discrepancy in residence times of inhibitor 5 and inhibitor 9 for the JAK2-inhibitor complex, PCA is implemented to the 3D positional coordinates obtained from the MD trajectories.^127–129^ PCA explains the variance in the dataset by transforming the MD trajectories into a set of orthogonal vectors or principal components representing characteristic molecular internal motions. The first PC shows the maximum variance in the data, followed by the second PC, and so on. Although the first PC is extremely useful in gaining insights into the system dynamics, the actual motion of the system is the combination of all the PCs. Figure 9a, and 9b shows the first PC obtained for the JAK2-inhibitor 5 and JAK2-inhibitor 9 complex, respectively. Figure 9a shows a greater domain movement around the binding region of the JAK2-inhibitor 5 complex. This motion may be attributed to a region of high instability around the binding site for inhibitor 5, leading to a lower residence time than inhibitor 9.

**Figure 9:**
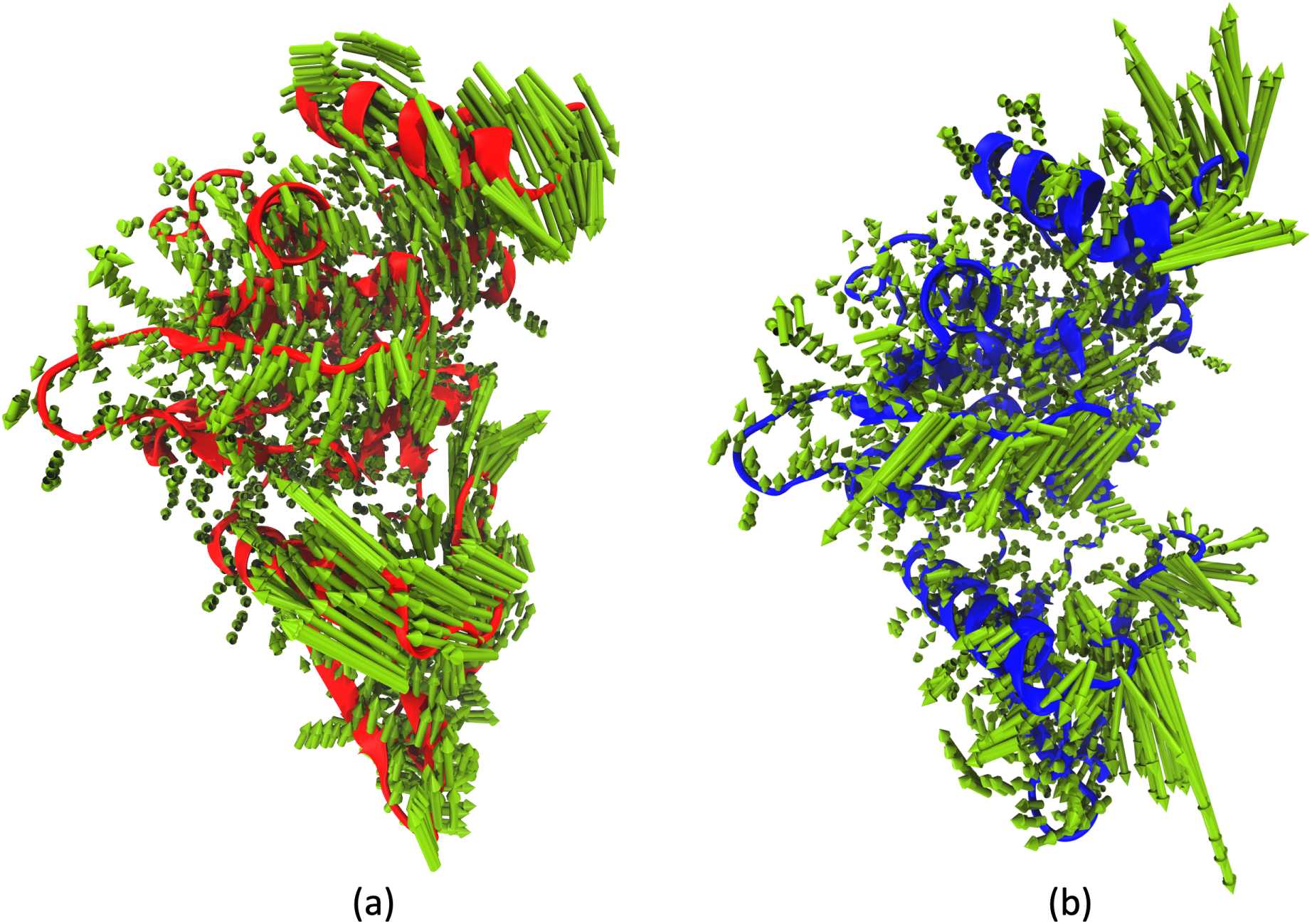
Principal Component Analysis for JAK2 inhibitor complexes from 2 *µ*s of MD simulation trajectory (a) First normal mode for JAK2-inhibitor 5 complex (47% of accounted variance) (b) First normal mode for JAK2-inhibitor 9 complex (46% of accounted variance)

#### 3.2.2 Selectivity of inhibitors towards JAK2 over JAK3

The inhibitors at the binding site of the JAK2 protein display higher residence times than the same series of inhibitors for the JAK3 protein. To corroborate these experimental findings, minimum average distance analysis is performed to obtain a detailed description of the binding pocket of the JAK-inhibitor complex. The minimum distance between any two atoms of the amino acid and the inhibitor averaged over the course of the 2 *µ*s trajectory for all the residues is calculated for the JAK-inhibitor complexes. Figure 10a represents the binding pocket of JAK2-inhibitor 9 complex, while Figure 10b represents the binding pocket of JAK3-inhibitor 9 complex. Interacting residues described in the figure are chosen with a cut-off distance of 4 Å. Supplementary Table S4 shows the list of interacting residues for inhibitor 9 in complex with JAK2 and JAK3 proteins. It is evident from Figure 10a and 10b that inhibitor 9 interacts with more residues of JAK2 over JAK3. It is also observed that the binding site occupies a larger volume, and the inhibitor is placed deeper in the binding pocket of JAK2, explaining the selectivity of the same towards JAK2 over JAK3. Interestingly for JAK3, it has been observed that the substituted fluorine atom in the heteroaryl C-ring in inhibitor 9 does not interact with the hydrogen of the *β*-carbon or any other heavy atom of the serine residue (Ser907).

**Figure 10:**
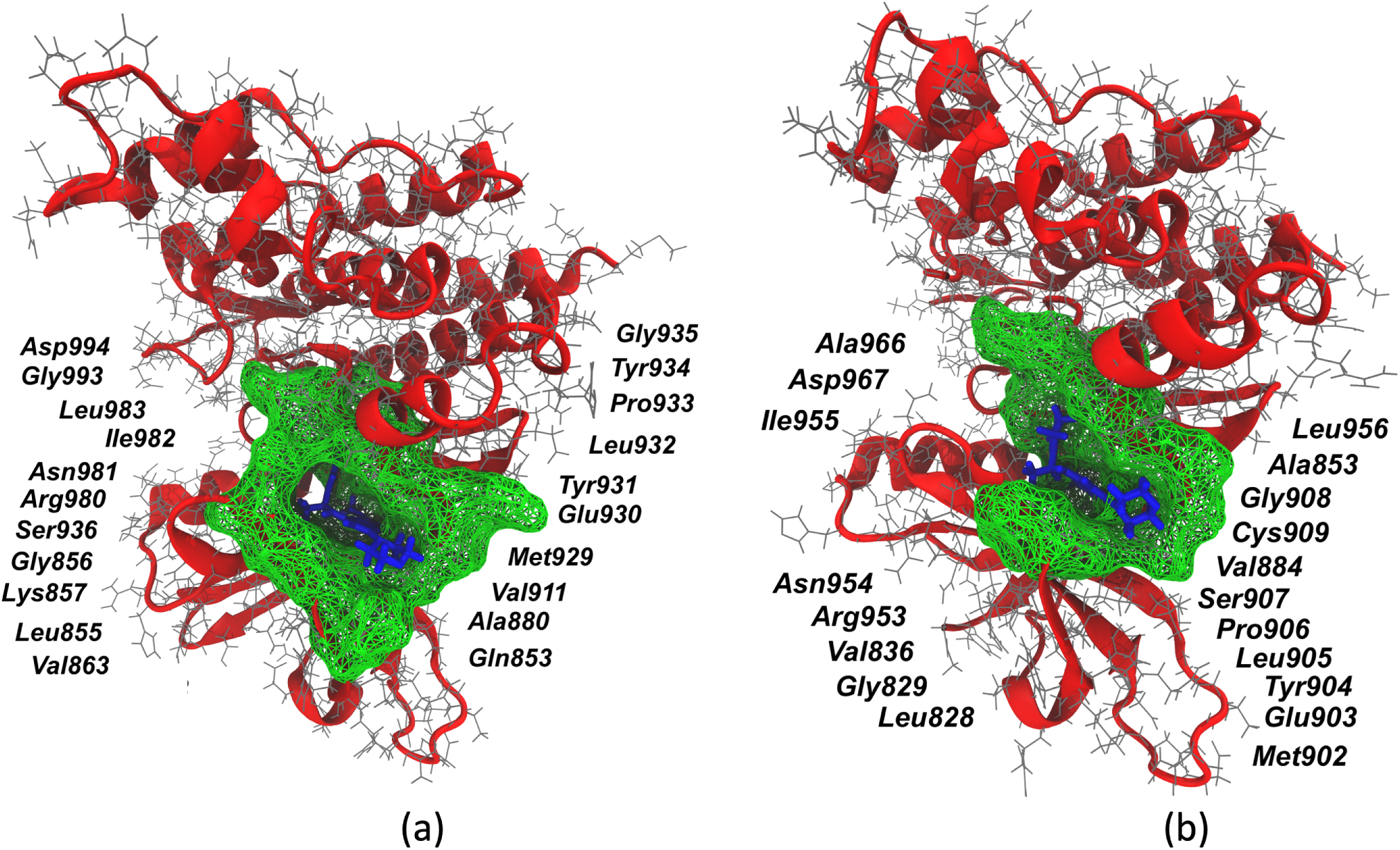
Binding site (green mesh) obtained from minimum average inhibitor-residue distances from three independent 2 *µ*s of MD simulation trajectories. (a) JAK2-inhibitor 9 complex (b) JAK3-inhibitor 9 complex

Root mean square fluctuations (RMSF) calculations are performed to identify important residues and domains associated with inhibitor binding and unbinding.^130^ A root mean squared (RMS) fit to the average structure is performed to obtain the fluctuations without rotations and translations, and a mass-weighted averaging of atomic fluctuations for each residue is carried out for the entire simulation trajectory. As demonstrated in Figure 11a, the binding site flanking residues for JAK2, namely Gly856, Lys857, Phe860, Gly861, Ser887, Glu889, Asp894, Arg897, Glu898, and Arg922, have lower RMSF and stabilize upon inhibitor 9 binding as compared to inhibitor 5 suggesting their role in stabilizing the receptor-inhibitor complex. Similarly, in JAK3 proteins, however, residue fluctuations are mostly similar, though only a few of the binding site flanking residues, such as Phe833, Gly834, Gln858, Gly861, Pro862, Asp863, Gln864, and Phe868, show a significant difference in fluctuations upon inhibitor 9 binding as compared to inhibitor 6 (Figure 11b). A higher number of residues in JAK2 contributing to the low fluctuations at the binding site may contribute to the selectivity of inhibitor 9 towards JAK2 over JAK3.

**Figure 11:**
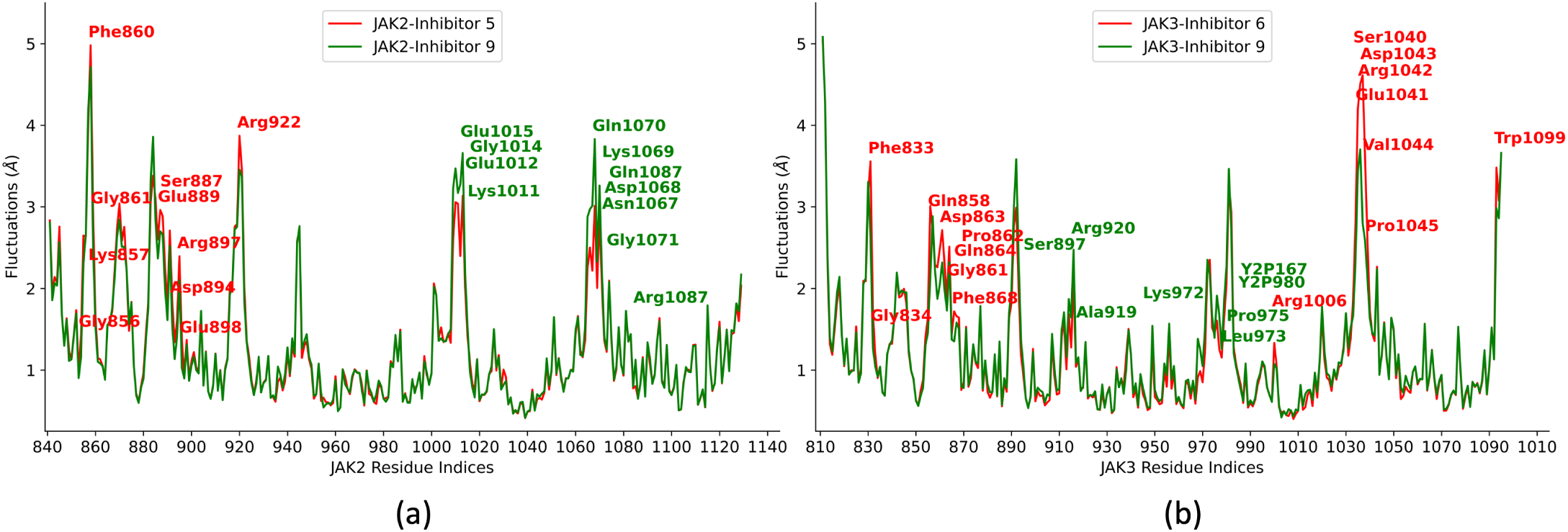
Residue fluctuation analysis for JAK2 and JAK3 inhibitor complexes obtained from three independent 2 *µ*s of MD simulation trajectories. (a) JAK2-inhibitor 5 vs. JAK2-inhibitor 9 complex (b) JAK3-inhibitor 6 vs. JAK3-inhibitor 9 complex

The binding pocket volume of the JAKs is a direct consequence of residues interacting with the inhibitor at the ATP binding site. These pocket volumes are complementary to the shape of the inhibitors as well. To compare the binding pockets of different inhibitors in JAK2 and JAK3 proteins, POVME, a tool to analyze binding pocket volumes, was utilized.^131, 132^ POVME provides a grid-based pocket representation of the inhibitor binding site. The pocket volumes are calculated with a grid spacing of 0.1 Å and a distance cut-off of 1.09 Å. Deep pocket volumes are observed for the JAK2 inhibitors where these inhibitors are tightly bound to the interacting residues. Supplementary Figure S1 shows a distinct difference in the binding pocket volumes for JAK2 vs. JAK3 proteins, where the volumes associated with inhibitors in the binding domain of JAK2 are significantly higher than those of JAK3.

Inhibitor-binding site distance analysis is performed for each receptor-inhibitor complex averaged over three independent MD simulation trajectories of 2 *µ*s each. From the starting structure of the zeroth milestone of each JAK-inhibitor complex, residues encompassing the inhibitor within a cut-off radius of 4 Å defined the binding site. The distance between the center of masses of the inhibitors and the *α*-C atoms of the binding site are used to calculate the inhibitor-binding site distance. It has been observed for all four inhibitors that the inhibitor-binding site distance in the case of JAK2-inhibitor complexes is less than that of the JAK3-inhibitor complexes (Supplementary Figure S2), suggesting strong binding of the inhibitors to the JAK2 protein.

JAK inhibitors target the JAK family of kinases and bind to the ATP-binding site of the kinase domain, thereby preventing the phosphorylation of downstream signaling proteins. In the case of JAK2 protein, the backbone amide and carbonyl groups (Leu855, Met929, and Leu932) interact with the phosphate groups of the ATP, forming multiple hydrogen bonds.^133, 134^ These interactions at the hinge region are of particular interest as they are conserved in the case of JAK2-inhibitor interactions (Figure 1). The inhibitors contain a heterocyclic core that mimics the adenine ring of the ATP to retain such interactions. Additionally, other interactions of these inhibitors with the kinase domain lead to the selectivity of these inhibitors over other kinases (Figures 8a and 8b).

## 4 Conclusion

The SEEKR2 milestoning method proved efficient in estimating the experimental residence times for different JAK-inhibitor complexes. The trend in residence times for the set of inhibitors for the JAK2 and JAK3 proteins is also conserved. It becomes evident from the SEEKR2 milestoning approach and the experiments that the series of inhibitors display an extended residence time and bind stronger to JAK2 than to JAK3. Among the inhibitors, inhibitor 9 displayed the highest residence time in the JAK2 protein. The results are further supported by MD simulations where important binding residues have lower distances from the inhibitor and lesser fluctuation in the JAK2-inhibitor 9 complex. In addition, the QM calculations show a higher electron density on the fluorine groups in the heteroaryl C-ring of inhibitor 9, strengthening the binding with JAK2 and JAK3 proteins resulting in the highest residence time among all the inhibitors. SEEKR2, thereby proves to be a valuable tool to predict the kinetics and thermodynamics of receptor-ligand binding and unbinding as it is user-friendly, requires minimum structural information of the system, is embarrassingly parallel, and requires a comparatively short simulation time to reach converged kinetic rates.

## Acknowledgement

The authors acknowledge Benjamin Jagger and Shiksha Dutta for insightful and helpful discussions. A.A.O. acknowledges the support of the Molecular Sciences Software Institute (MolSSI) fellowship under NSF grant OAC-1547580. R.E.A. acknowledges support from NSF Extreme Science and Engineering Discovery Environment (XSEDE) CHE060063 and NIH GM132826. All simulations were performed using the Triton Shared Computing Cluster (TSCC) and Popeye computing cluster at the San Diego Supercomputing Center (SDSC).

## Supporting Information Available

Supplementary information includes the list of interacting residues of JAKs with inhibitors, the pocket volume analysis, and the inhibitor-binding site analysis for different JAK-inhibitor complexes.

## Data and Software Availability

The SEEKR2 project is available at https://github.com/seekrcentral/seekr2. The structures of JAK2 and JAK3 proteins complexed with the inhibitors, analysis scripts, and scripts for system preparations for SEEKR2 simulations are available at https://github.com/anandojha/kinase_SEEKR. The data for this study can be found at https://doi.org/10.6075/J01Z44MN.

## TOC Graphic

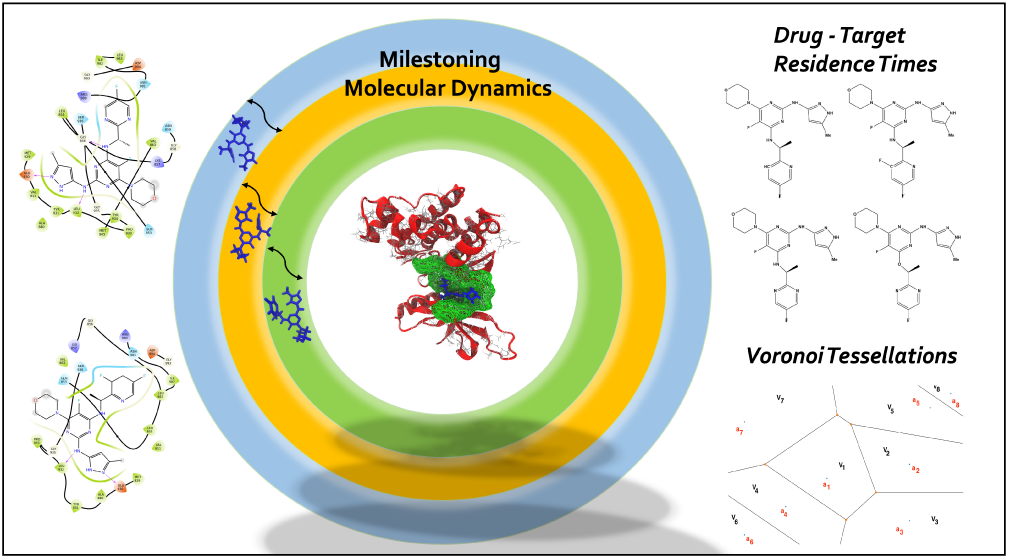

