## Supplementary Information File for "Selectivity and ranking of tight-binding JAK-STAT inhibitors using Markovian milestoning with Voronoi tessellations"

Table S1: JAK2 and JAK3 residues within a cut-off distance of 3 Å of the ligand in the bound state (defined as binding site residues).

| JAK-inhibitor complex | Interacting Residues |
|---|---|
| JAK2-inhibitor 5 | Leu855, Gly856, Lys857, Val863, Ala880, Val911, Met929, Glu930, Tyr931, Leu932, Pro933, Gly935, Ser936, Arg980, Asn981, Leu983, Gly993 |
| JAK2-inhibitor 6 | Leu855, Gly856, Lys857, Val863, Ala880, Met929, Glu930, Tyr931, Leu932, Pro933, Gly935, Ser936, Arg980, Asn981, Ile982, Leu983, Gly993 |
| JAK2-inhibitor 7 | Gln853, Leu855, Val863, Ala880, Met929, Glu930, Tyr931, Leu932, Pro933, Gly935, Ser936, Leu983, Gly993, Asp994 |
| JAK2-inhibitor 9 | Leu855, Gly856, Lys857, Val863, Ala880, Val911, Met929, Glu930, Tyr931, Leu932, Pro933, Tyr934, Gly935, Ser936, Leu983, Gly993 |
| JAK3-inhibitor 5 | Leu828, Gly829, Val836, Ala853, Val884, Met902, Glu903, Tyr904, Leu905, Pro906, Gly908, Cys909, Arg953, Asn954, Leu956, Ala966 |
| JAK3-inhibitor 6 | Leu828, Val836, Ala853, Val884, Met902, Glu903, Tyr904, Leu905, Pro906, Gly908, Cys909, Arg916, Arg953, Asn954, Leu956, Ala966, Asp967 |
| JAK3-inhibitor 7 | Leu828, Gly829, Val836, Ala853, Val884, Met902, Glu903, Tyr904, Leu905, Pro906, Gly908, Cys909, Arg953, Asn954, Leu956, Ala966, Asp967 |
| JAK3-inhibitor 9 | Leu828, Gly829, Val836, Ala853, Val884, Met902, Glu903, Tyr904, Leu905, Pro906, Gly908, Cys909, Arg953, Asn954, Leu956, Asp967 |

Table S2: Experimentally determined vs. the SEEKR2 calculated residence times of inhibitors with the kinase domain of the JAK2 and JAK3 proteins. The SEEKR2 residence times and error estimates are the mean of each of the three SEEKR2 simulations.

| JAK-inhibitor complex | Experimental Residence Time (hrs.) | SEEKR2 Residence Time (hrs.) |
|---|---|---|
| JAK2-inhibitor 5 | 2.68 | $1.70 \pm 0.004$ |
| JAK2-inhibitor 6 | 6.65 | $6.79 \pm 0.012$ |
| JAK2-inhibitor 7 | 3.20 | $2.31 \pm 0.007$ |
| JAK2-inhibitor 9 | 11.07 | $10.96 \pm 0.096$ |
| JAK3-inhibitor 5 | 0.98 | $0.54 \pm 0.003$ |
| JAK3-inhibitor 6 | 0.78 | $0.19 \pm 0.001$ |
| JAK3-inhibitor 7 | 1.30 | $1.12 \pm 0.006$ |
| JAK3-inhibitor 9 | 3.15 | $2.53 \pm 0.011$ |

Table S3: Unpaired t-test to measure the statistical significance of the difference between the SEEKR2 calculated residence times of inhibitors with the kinase domain of the JAK2 and JAK3 proteins with the experimentally determined residence times.

| JAK-inhibitor complex | SEEKR2 Mean Residence Time (hrs.) | Experimental Residence Time (hrs.) | t-ratio | p-value | Statistically Significant Difference ($\alpha$=0.05) |
|---|---|---|---|---|---|
| JAK2-inhibitor 5 | 1.702 | 2.68 | 1.132 | 0.375148 | No |
| JAK2-inhibitor 6 | 6.792 | 6.65 | 0.1065 | 0.924872 | No |
| JAK2-inhibitor 7 | 2.306 | 3.20 | 1.082 | 0.392391 | No |
| JAK2-inhibitor 9 | 10.960 | 11.07 | 0.0177 | 0.987493 | No |
| JAK3-inhibitor 5 | 0.5379 | 0.98 | 1.26 | 0.334802 | No |
| JAK3-inhibitor 6 | 0.1872 | 0.78 | 3.597 | 0.069358 | No |
| JAK3-inhibitor 7 | 1.116 | 1.30 | 0.119 | 0.916129 | No |
| JAK3-inhibitor 9 | 2.535 | 3.15 | 0.3385 | 0.767246 | No |

Table S4: Residues of JAK proteins interacting with inhibitor 9 obtained from the minimum average distance analysis within a cut-off distance of 4 Å averaged over three independent 2 $\mu$s MD simulation trajectories.

| JAK-inhibitor complex | Interacting Residues |
|---|---|
| JAK2-inhibitor 9 | Gln853, Leu855, Gly856, Lys857, Val863, Ala880, Val911, Met929, Glu930, Tyr931, Leu932, Pro933, Tyr934, Gly935, Ser936, Arg980, Asn981, Ile982, Leu983, Gly993, Asp994 |
| JAK3-inhibitor 9 | Leu828, Gly829, Val836, Ala853, Val884, Met902, Glu903, Tyr904, Leu905, Pro906, Ser907, Gly908, Cys909, Arg953, Asn954, Ile955, Leu956, Ala966, Asp967 |

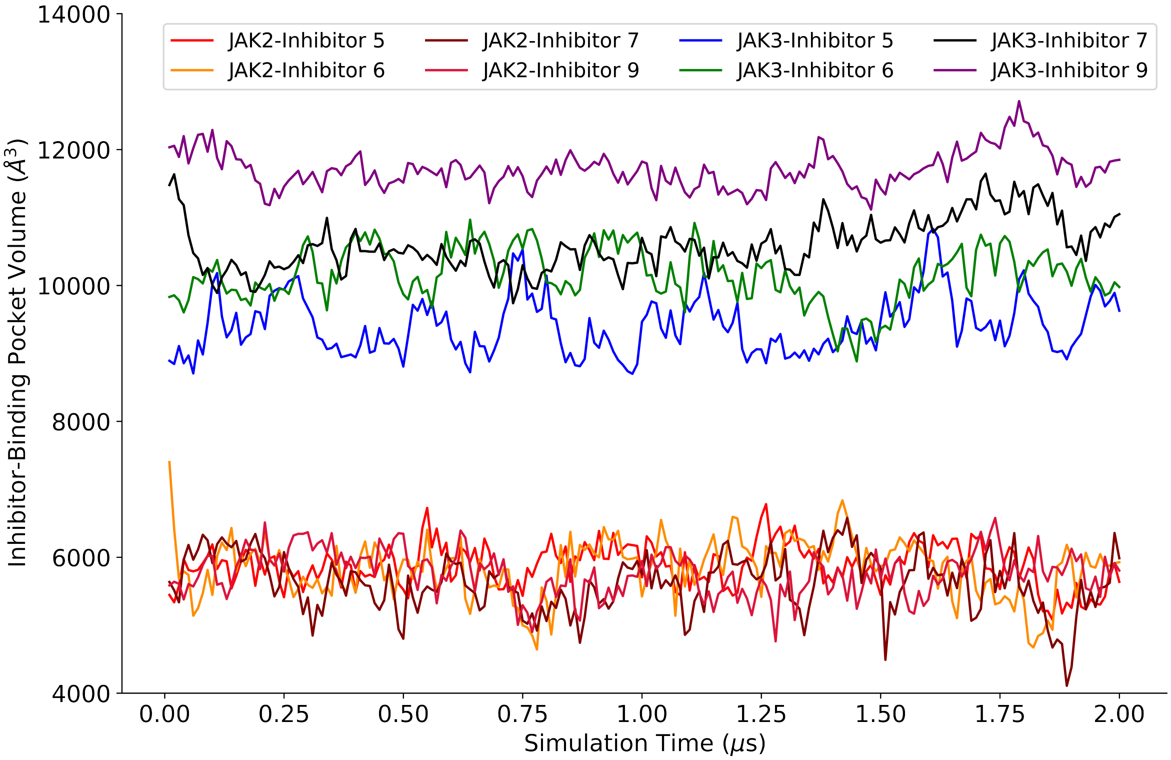


Figure S1: Pocket volume analysis for all the JAK2 and JAK3 inhibitor complexes.

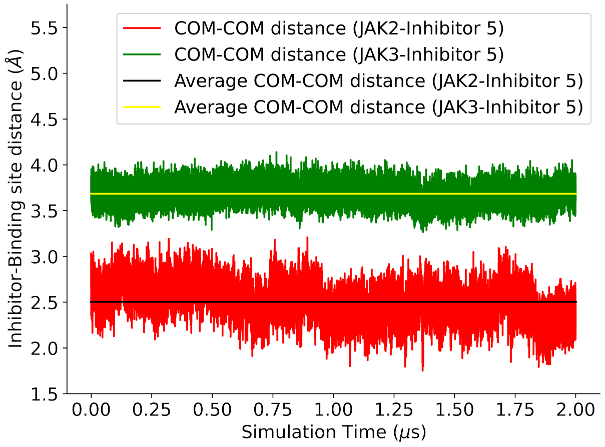


(a)

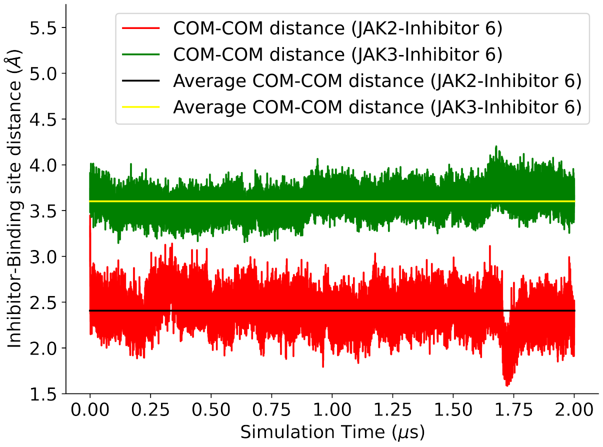


(b)

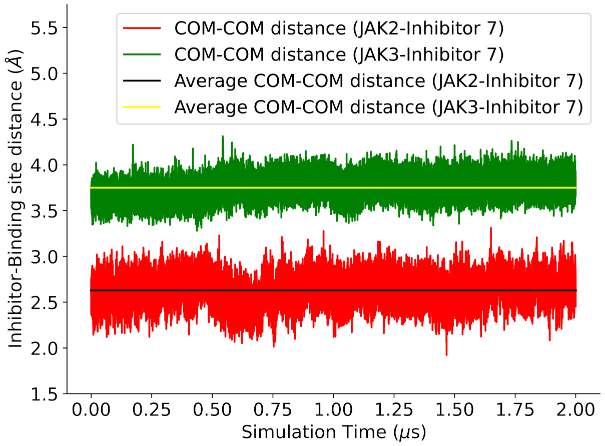


(c)

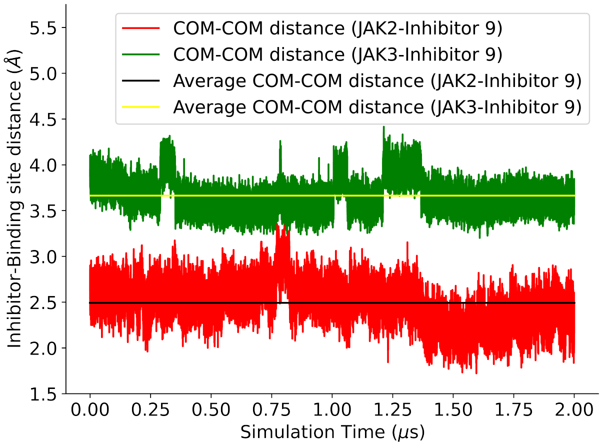


(d)

Figure S2: Inhibitor-Binding site distance analysis for JAK2 and JAK3 inhibitor complexes from three independent 2 $\mu$s MD simulation trajectories. The distance between the center of masses of the inhibitors and the $\alpha$-C atoms of the binding site are used to calculate the inhibitor-binding site distance. (b) JAK2-inhibitor 5 vs. JAK3-inhibitor 5 complex (c) JAK2-inhibitor 6 vs. JAK3-inhibitor 6 complex (d) JAK2-inhibitor 7 vs. JAK3-inhibitor 7 complex (e) JAK2-inhibitor 9 vs. JAK3-inhibitor 9 complex
